# Spontaneous generation of face recognition in untrained deep neural networks

**DOI:** 10.1101/857466

**Authors:** Seungdae Baek, Min Song, Jaeson Jang, Gwangsu Kim, Se-Bum Paik

**Affiliations:** Department of Bio and Brain Engineering, Korea Advanced Institute of Science and Technology, Daejeon 34141, Republic of Korea; Program of Brain and Cognitive Engineering, Korea Advanced Institute of Science and Technology, Daejeon 34141, Republic of Korea; Department of Physics, Korea Advanced Institute of Science and Technology, Daejeon 34141, Republic of Korea

## Abstract

Face-selective neurons are observed in the primate visual pathway and are considered the basis of facial recognition in the brain. However, it is debated whether this neuronal selectivity can arise spontaneously, or requires training from visual experience. Here, we show that face-selective neurons arise spontaneously in random feedforward networks in the absence of learning. Using biologically inspired deep neural networks, we found that face-selective neurons arise under three different network conditions: one trained using non-face natural images, one randomized after being trained, and one never trained. We confirmed that spontaneously emerged face-selective neurons show the biological view-point-invariant characteristics observed in monkeys. Such neurons suddenly vanished when feedforward weight variation declined to a certain level. Our results suggest that innate face-selectivity originates from statistical variation of the feedforward projections in hierarchical neural networks.

## Introduction

The ability to identify and recognize faces is a crucial function in visual-priority social animals such as humans and other primates, and is thought to originate from neuronal tuning at a single or multi-neuronal level. Neurons that selectively respond to faces (face-selective neurons) are observed to occur in the inferior temporal cortex (IT)^1–6^, superior temporal sulcus (STS)^7–10^, and fusiform face area (FFA)^11–15^ in the primate brain (**Fig. 1A**). Several contradictory observations on the origin of face-selective neurons in infant animals have been reported, raising two different scenarios for the development of this intriguing functional tuning.

**Figure 1.**
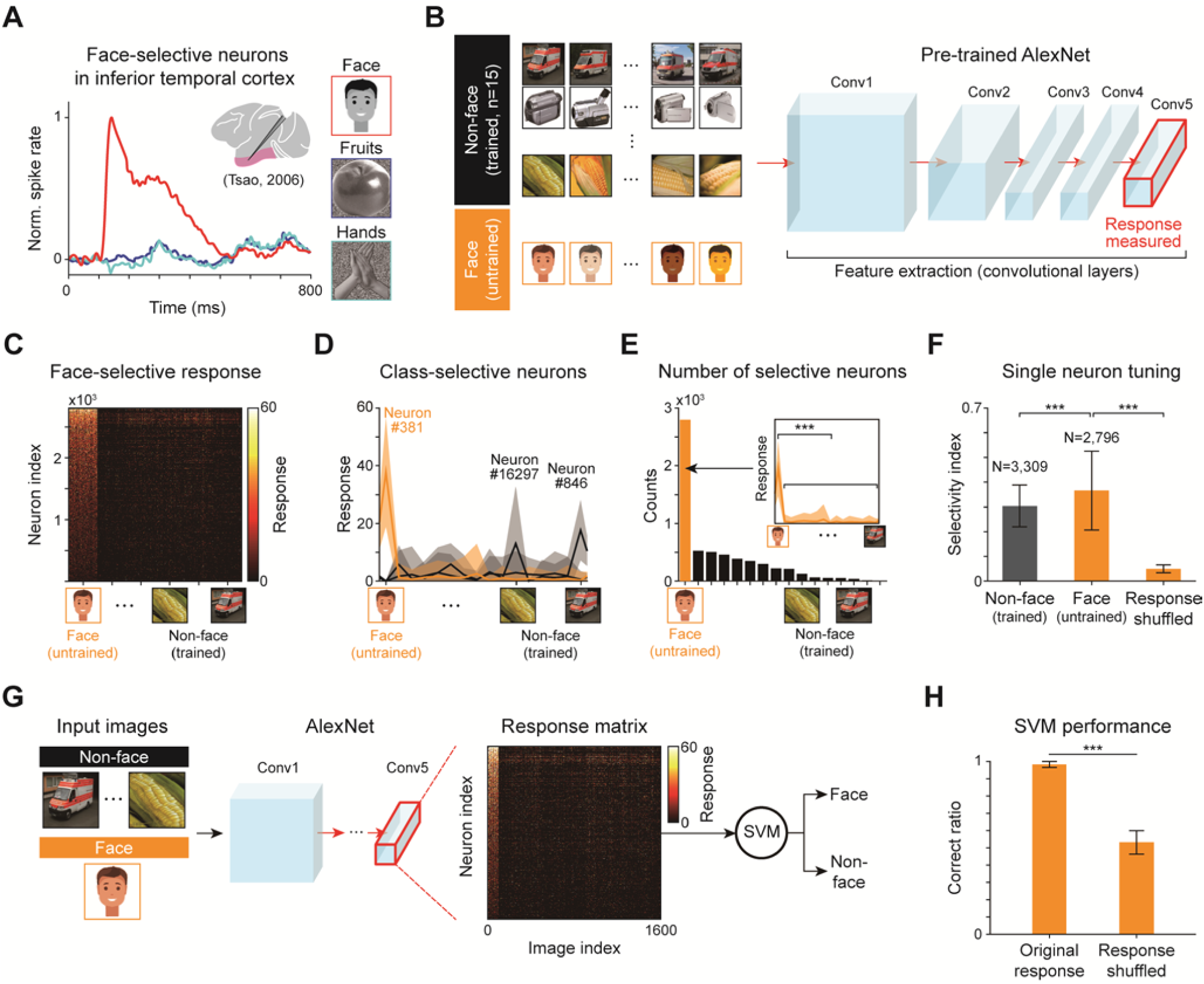
Emergence of face-selectivity in networks trained for non-face natural image classification. **A** Face-selective neurons observed in monkey experiments^1^. Note that, according to the policy of bioRxiv that avoid the inclusion of photographs and any other identifying information of people, here we show the illustration of faces instead of actual face pictures we used throuout this manuscript. **B** (Left) Non-face natural images in ImageNet^50^, which is trained in AlexNet^27^, and face image not trained in the network. (Right) Schematic architecture of the pre-trained AlexNet. **C** Face-selective responses of individual face-selective network neurons. **D** Example for tuning curves of individual neurons selective for different image classes. The shaded area represents the standard deviation of the response distribution obtained from 100 different images of the selective class. **E** Distribution of the number of neurons for each preferred image class. The number of selective neurons for each class is sorted from left to right in decreasing order. (Inset) the selective neuron is defined as the neuron which show significantly higher response to preferred image class than any other class (***p < 0.01, Mann–Whitney U test). **F** Selectivity of individual face-selective and non-face-selective neurons defined as the probability that a neuron shows a maximum response to an image of their selective class. The error bar represents the standard deviation of the selectivity index for each condition. **G** Design on the face classification task and SVM classifier using the responses of AlexNet. **H** Performance on the Face classification task using the original response and permuted response matrix. The error bar represents the standard deviation of the performance of 20 permuted response matrixes.

The first scenario is that visual experience develops face-selective neurons. A study using functional magnetic resonance imaging (fMRI) to examine FFA in monkeys reported that the category of selective neuronal activity observed, depends greatly on what a subject had experienced in its lifetime^16^. Another fMRI study of IT in monkeys reported that robust tuning of face-selective neurons is not observed until one year after birth^5^ and that face-selectivity relies on experience during the early infant years. Furthermore, it was reported that monkeys raised without face exposure did not develop normal face-selective domains^17^. These results suggest that face-selective neurons are developed from training with visual experience.

However, the other view suggests that face-selectivity can innately arise without visual experience^18–21^. It was observed that human infants behaviorally prefer to look face-like objects rather than non-face ones^22–24^, implying that face-encoding units may already exist in infants. It was also reported that adult humans with no visual experience have category-selective domains including face, in the ventral visual cortex^19^. In addition, a recent fMRI study of infant animals reported that face-selective neurons are observed with movie stimulus, but not observed with static image inputs^5^. Furthermore, the spatial organization of such early face-selective regions appeared similar to that observed in adults. These results, altogether, imply that face-selective neurons might arise innately without visual experience, in contradiction to the first scenario.

These contradictory results were probably due to limitations in the control of the experimental conditions. For example, it is impossible to control the amount of visual experience for a particular category, such as face, in individual subjects. Even if the subjects are visually deprived so that they are prevented from having visual experience, the portion of category-selective neurons and their degree of tuning may vary across subjects and cannot easily be predicted. These various factors make it difficult to investigate the developmental mechanism of face-selectivity in the brain.

A model study using biologically-inspired artificial neural networks, such as deep neural networks (DNNs)^25^, might offer an alternative approach in this case^26–28^. Recently, model studies with DNNs have successfully provided insight into the underlying mechanisms of brain functions, particularly with regard to the development of various functions for visual perception^29–31^.

Herein, we show that face-selective neurons can spontaneously arise in completely untrained neural networks. Using DNNs reproducing the structure of the ventral stream of the visual cortex, we found that face-selective neurons arise under three different network conditions: one trained for non-face natural images, one randomized after being trained, and one randomly initialized and never trained. We observed that spontaneously emerged face-selective neurons show the biological characteristics of view-point invariance observed in IT of monkeys. From further investigation, we found that face-selective neurons can emerge from the statistical variation of feedforward projection weights in the network. We also found that face-tuning vanishes when the feedforward variation declines. Our findings suggest that innate face recognition may originate from face-selective neurons that emerge spontaneously from the early development of random feedforward wiring in the visual pathways.

## Results

### Emergence of face-selective neurons in networks trained for non-facial natural images

To investigate the development of face-selective neurons (**Fig. 1A**) in a biologically inspired deep neural network (DNN) model, we implemented an adapted version of AlexNet^27^ (**Fig. 1B**). A standard AlexNet model is composed of five convolutional layers (feature extraction network) and three fully-connected layers (classification network), which together reproduce the structure of the ventral stream of the visual pathway. To investigate the selective response of individual neurons rather than the performance of a trained system, we discarded the classification layers and examined neuronal activity in the final layer (conv5) of the feature extraction network.

We first tested whether face-selective neurons could arise in a network trained to only non-face natural images. Using an AlexNet pre-trained with non-face natural images (ImageNet database N = 1,000 classes, see Methods for details), we measured neural responses to the stimulus image sets of face (untrained) and non-face (N = 15 selected trained classes) (**Supplementary Fig. S1**). We found that face-selective neurons were observed in this condition (**Fig. 1C** and **D**). We found 2,796 face-selective neurons (out of 43,264 neurons in the final layer) that showed significantly higher response to face images compared to non-face images (**Fig. 1E**, inset, p < 0.01, Mann–Whitney U test). Furthermore, we found that the number of face-selective neurons that emerged was much greater than that of neurons selective for each trained class of non-face objects (**Fig. 1E**). To quantify the degree of tuning in individual neurons, we defined the selectivity index as the probability that a neuron generates a maximum response to the preferred class of images^32^ (see Methods for details). Face-selective neurons showed very sharp tuning to face images such that their average selectivity index was significantly higher than that of a control obtained from shuffled responses, and even slightly higher than that of neurons tuned to the trained classes (**Fig. 1F, \*\*\***p < 0.001, face class: 0.36 ± 0.16, non-face class average: 0.30 ± 0.08, response of neurons shuffled: 0.04 ± 0.02, Mann–Whitney U test).

Next, we tested whether the selective response of these neurons could provide sufficient information for successful performance in a face classification task (**Fig. 1G**). In this task, face (N = 60) or non-face (N = 60) images were randomly presented to the network, and the observed neuronal response of the final layer was used to train a support vector machine (SVM) to classify whether the given image was a face or not (see Methods for details). We confirmed that the network successfully performed the task from the activity of face-selective neurons. The measured correct performance rate of the network was found to be 0.98 ± 0.02 (for N = 60 test images). This is significantly higher than with the control of which the responses of the neurons were shuffled across two presented images (0.53±0.07, **Fig. 1H**, ***p < 0.001, Mann–Whitney U test). This result implies that the face-selective neurons that spontaneously emerged in the AlexNet trained to non-face natural images can provide the network with the capability to distinguish a face.

### Spontaneous emergence of face-selective neurons in untrained networks

Next, to validate whether face-selective neurons indeed arise from the training process for object classification, we devised an untrained AlexNet (permuted AlexNet) by randomly permuting the weights of kernels in each convolutional layer of the pre-trained network (**Fig. 2A**). Surprisingly, even though the network was never trained with visual stimuli after the randomization step, face-selective neurons (**Fig. 2B**, e.g., neuron #13527) and non-face (neurons #40309 and #11651) classes were observed in the permuted AlexNet. Similar to pre-trained network, the number of face-selective neurons in the permuted network was significantly greater than the number of neurons responsive to non-face objects (**Fig. 2C**, ***p < 0.001, face-selective = 1,601 ± 275, non-face-selective: 398 ± 27, Mann–Whitney U test). The observed face-selective neurons showed sharp tuning curves similar to the ones observed in the pre-trained network, and their average selectivity index was significantly higher than that of shuffled responses. Furthermore, we also found that the average selectivity index of face-selective neurons appeared slightly higher than that of the neurons responsive to non-face objects (**Fig. 2D**, ***p < 0.001, Mann–Whitney U test).

**Figure 2.**
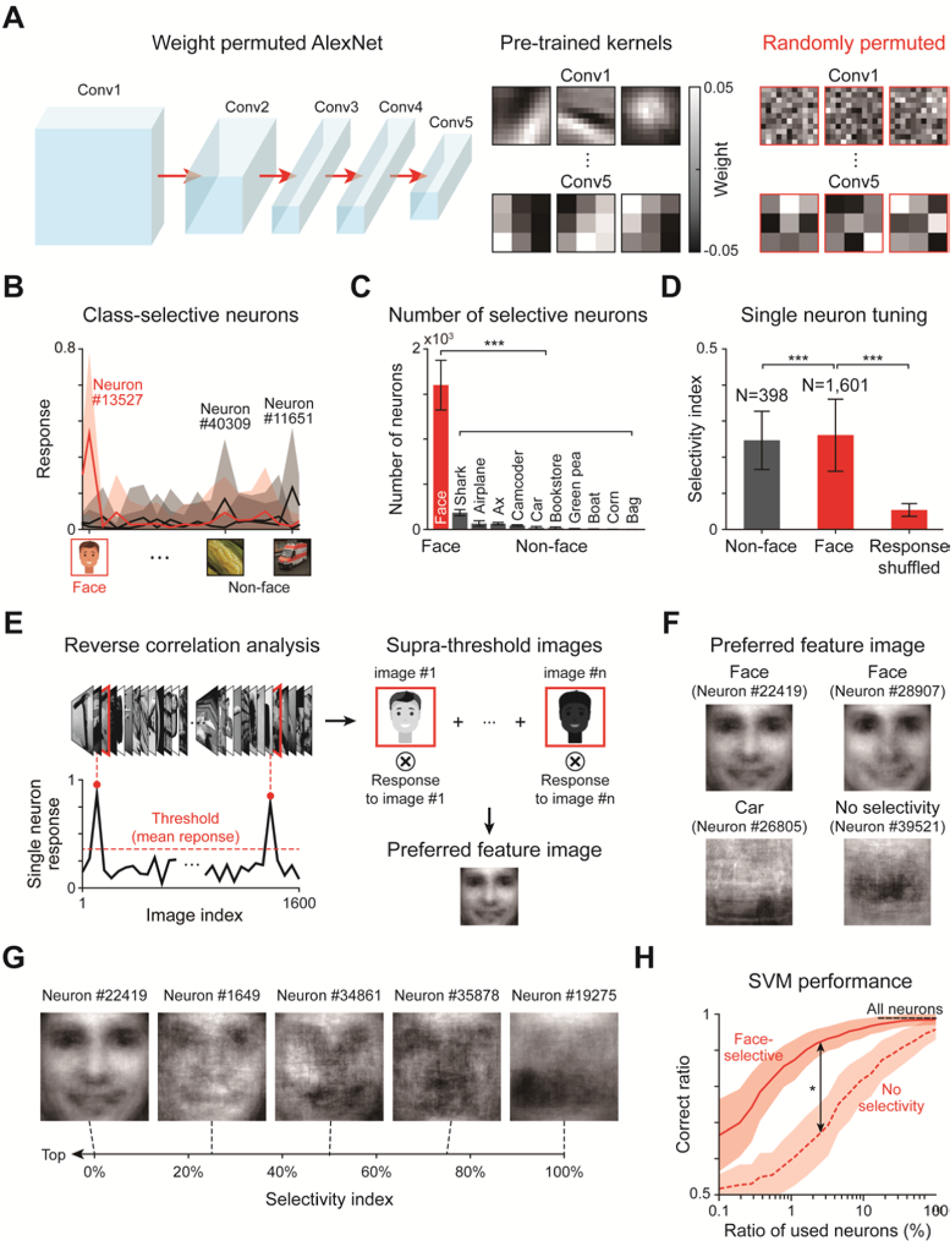
Spontaneous emergence of face-selectivity in untrained networks. **A** The untrained AlexNet was devised by randomly permuting the weights in each convolutional layer. **B** Examples of tuning curves for individual face-selective network neurons. **C** Number of neurons responsive to each image class, sorted from left to right in decreasing order. Error bars represent the standard deviation on 100 permuted networks. **D** Selectivity of individual face-selective and non-face-selective neurons. The error bar represents the standard deviation of all the selectivity indices on 100 permuted networks. **E** Measurement of preferred feature images of target neuron in conv5 from reverse correlation analysis. Stimuli inducing responses higher than a threshold (average response of all neurons to all images) were selected, weighted by corresponding responses and added. **F** The preferred feature images of sample neurons that are face-selective, car-selective, and with no selectivity. **G** Preferred feature images of face-selective neurons with various selectivity indices. **H** Performance of the face classification task using (1) all neurons (N_all_ = 43,264), (2) only face-selective neurons (100% face-selective neurons: N_face_ = 1,601), (3) neurons with no selectivity. The shaded area represents the standard deviation of performance on 100 trials.

To characterize qualitatively the response of these face-selective neurons, we reconstructed the preferred feature images of individual neurons using the reverse correlation method (**Fig. 2E**). For this, we presented 1,600 natural images including face and 15 non-face classes to the permuted network, and then selected images that induced a response above the mean response of all neurons to all images (See Methods). By adding supra-threshold images weighted by corresponding neural response, we obtained preferred feature images of three different neuron groups: (1) face-selective neurons, (2) neurons selective for non-face objects, (3) neurons with no selectivity. In face-selective neurons, face-like shapes including components such as eyes, nose, and mouth were observed in preferred feature images (**Fig. 2F**). In some cases of neurons responsive to non-face objects, partial silhouettes such as a part of car were observed, but not clearly visible as in case of face. No noticeable shape was detected in neurons with no selectivity, as expected. Furthermore, we confirmed that face-like shapes in the preferred feature image of face-selective neurons were more clearly noticeable in neurons of higher selectivity index (**Fig. 2G**).

To test whether face-selective neurons that spontaneously emerged in the untrained network could also enable the network to classify face images among other objects, we repeated the face classification task with a support vector machine (SVM) by changing the number of face-selective neurons used for SVM. We first found that the SVM trained only with face-selective neurons (N_face_ = 1,601) showed performance comparable with that using all neurons (N_all_ = 43,264) in the final layer (**Fig. 2H**, Face-neurons: 0.98 ± 0.02, All neurons: 0.99 ± 0.01). This implies that face-selective neurons can provide the network with the capability to distinguish a face. To confirm further that the response of face-selective neurons enabled this result, we compared the classification performance of an SVM using the same number (N = 1,601) of randomly sampled neurons with no class-selectivity. We confirmed that the SVM trained with only face-selective neurons shows noticeably better performance than that with neurons without class-selectivity, as the number of neurons used in each condition was varied from N = 2 (0.1% of the total face-selective neurons) to N = 1,601 (100%) (*p < 0.05, Kolmogorov-Smirnov test). This result implies that the tuned activity of face-selective neurons can induce innate face recognition.

### View-point invariant response of face-selective neurons

In previous reports from the fMRI study on monkeys, it was observed that the face-selective neurons in the inferior temporal cortex (IT) show responses invariant to diverse angles of the face images, a condition called view-point invariance^6^ (**Fig. 3A**). It was also observed that neurons show an increasing trend of invariance from middle lateral (ML) to anterior medial (AM) IT, as it goes to the higher hierarchy in IT (**Fig. 3B**). In subsequent analysis, we found that the face-selective neurons that spontaneously emerged in our untrained networks, reproduced view-point invariant profiles, consistent with that observed in biological data^6^.

**Figure 3.**
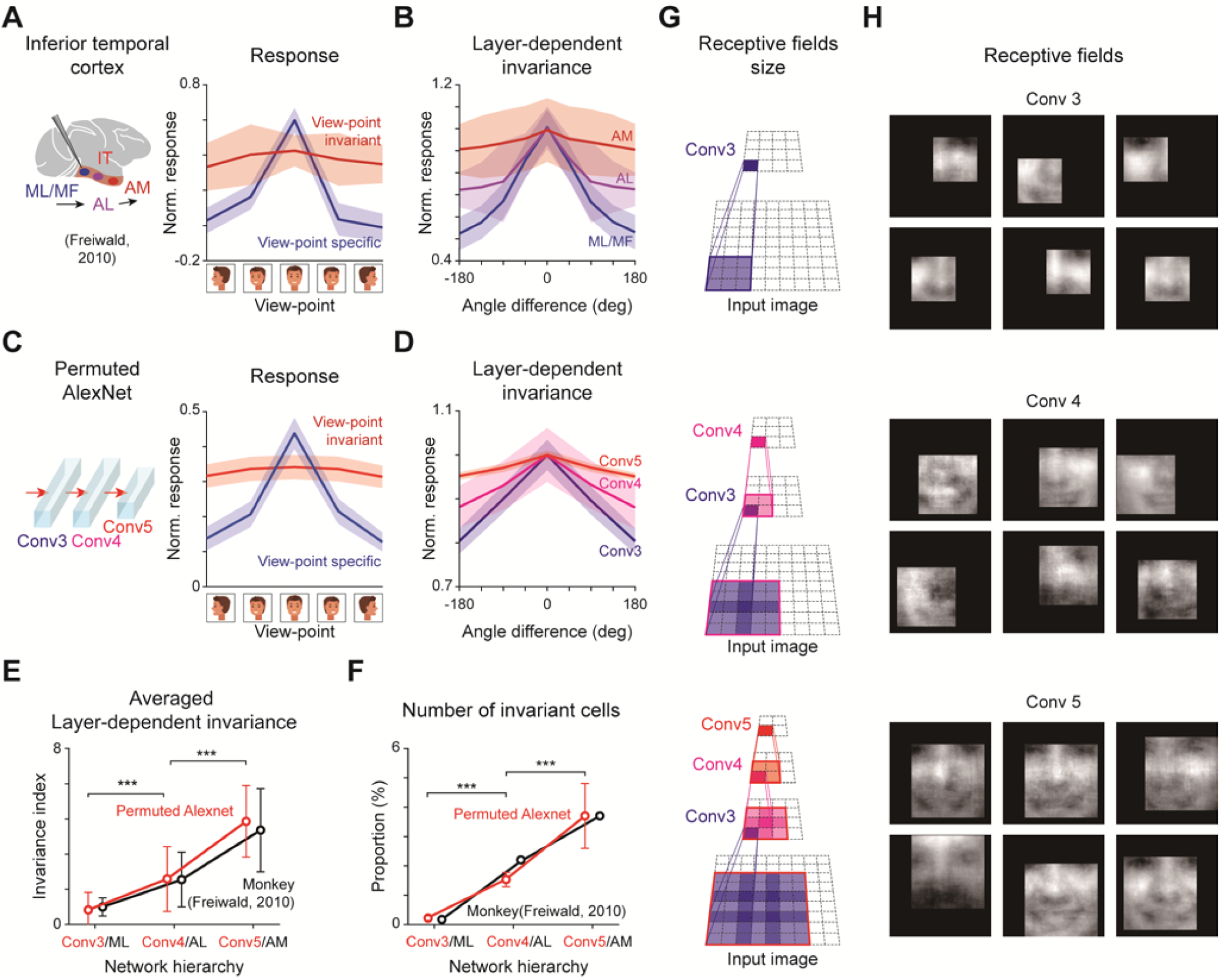
View-point invariant characteristics of face-selective network neurons. **A** View-invariant and view-specific response of face-selective neurons in monkey IT^6^. **B** Average tuning curves of face-selective neurons in each layer, which reveals increasing view-invariant characteristics along the IT hierarchy. **C** View-invariant and view-specific response of face-selective neurons in permuted AlexNet. The shaded area represents the standard deviation of the response on 100 permuted networks. **D** Average tuning curves of face-selective neurons in each layer. **E** Increasing invariance index over layer in both permuted AlexNet and monkey IT. The error bar represents the standard deviation of invariance indices for the three layers of 100 permuted networks. **F** The number of view-invariant neurons increases over layer in both permuted AlexNet and monkey IT. **G** The size of the neuronal receptive field in each convolutional layer was calculated from the deconvolution process. **H** The receptive field was obtained by cropping the net input area of the target neuron on the preferred feature image.

To investigate the view-point invariant characteristic of face-selective neurons, we measured the response of the permuted AlexNet while artificially generated face images (FaceGen Modeler software, singular inversions) from different angles were provided to the network (**Fig. 3C, Supplementary Fig. S2**, see Methods for details). We found that view-point invariant responses of face-selective neurons were observed, and that their level of invariance was increased along the network hierarchy in the permuted AlexNet, similar to that in monkey IT (**Fig. 3D**). To quantify these invariant characteristics, we introduced an invariance index of neurons, defined as the inverse of the response variance across different view angles. As a result, we found that higher layers (conv4 and 5) show relatively higher invariance than that in lower layers (conv3), consistent with observed monkey data^6^ (**Fig. 3E**, ***p < 0.001, Mann–Whitney U test). In addition, the number of view-point invariant neurons increased in higher layers in the network hierarchy, also similar to the condition observed in monkeys^6^ (**Fig. 3F**, ***p < 0.001, Mann– Whitney U test). These findings show that face-selective neurons spontaneously generated in untrained networks have biologically realistic characteristics similar to those observed in monkey IT, not only in single cells but also at population and inter-layer levels.

To examine the origin of such invariant characteristics, we examined the receptive fields of face-selective neurons, further considering the location and size of receptive fields in each layer. We backtracked the convolutional feedforward inputs and calculated the correspondent receptive fields of each neuron (**Fig. 3G**, top). As a result, we found that face-selective neurons in the lower layers detect only local compartments of a face, such as eyes, nose, and mouth, the shape of which sensitively varies by face angle. On the other hand, neurons in higher layers were observed to integrate local components and detect the features of the whole face, the profile of which is more consistent with variation of face angle (**Fig. 3G**, bottom). These results are consistent with the observed view-specific characteristics of neurons in the lower layers and the view-invariant characteristics of neurons in the higher layers.

### Face-selectivity arise from diversity of convolutional weight

To examine the origin of face-selectivity in untrained neural networks, we implemented a randomly initialized network (randomized AlexNet) where values in each weight kernel were randomly drawn from a Gaussian distribution that fit the weight distribution of the pre-trained state (**Fig. 4A**). Here, the variation of weights in the feedforward kernels could be controlled by modulating the width of the Gaussian (*σ*, a standard deviation of the weight distribution). Using this network, we investigated whether face-selective neurons could spontaneously arise from the weight variation of random feedforward networks.

**Figure 4.**
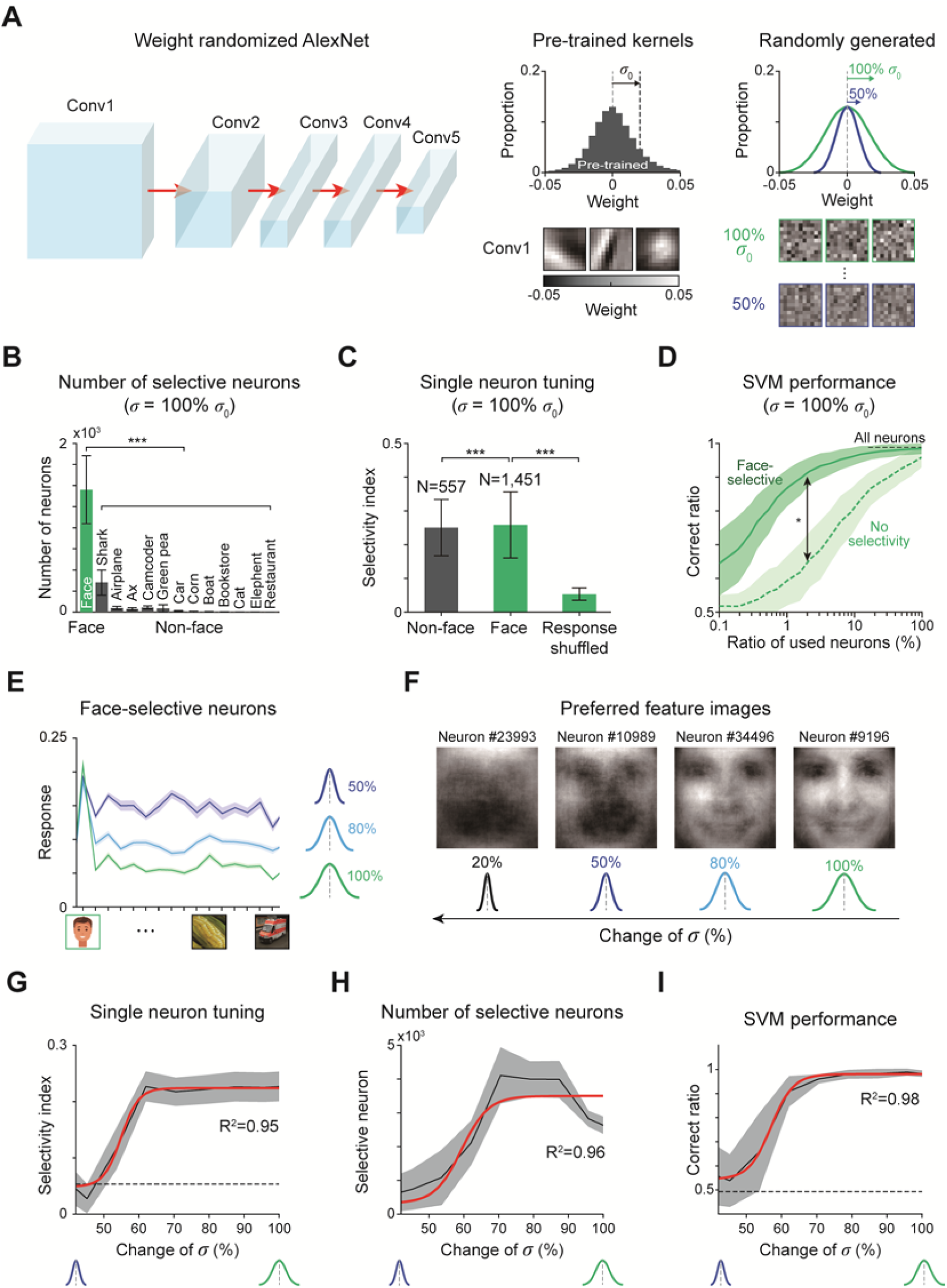
Face-selectivity induced by variation in convolutional weightsy. **A** Untrained AlexNet was generated, where values in each weight kernel were randomly sampled from a Gaussian distribution that fit the weight distribution of the pre-trained state. **B** Number of neurons responsive to each image class, sorted from left to right in decreasing order. Error bars represent the standard deviation on 100 permuted networks. **C** Selectivity of individual face-selective and non-face-selective neurons. The error bar represents the standard deviation of all selectivity indices on 100 randomly initialized networks. **D** Performance of face classification task in using face-selective neurons (100% face-selective neurons: N_face_ = 1,451), and the same number of neurons responsive to non-face objects. The shaded area represents the standard deviation of performance on 100 trials. **E** Average of all tuning curves for face-selective neurons across different levels of weight variation. Tuning becomes broader as the weight variation is reduced. **F** The disruption of receptive fields as the weight variation is reduced. The preferred feature images of face-selective neurons with the highest selectivity index was shown for each network with different weight variation. **G** The selectivity index of face-selective neurons where the weight variation was changed from 42% to 100% of original value. The gray shaded area represents the standard deviation on 100 trials. Red solid lines indicate fitting for the sigmoid function: *a*/(1 + *e*^−*bx*+*c*^) + *d*) (G: R^2^ of fit for the sigmoid function = 0.95, *p* < 10^−4^; H: R^2^ of fit for the sigmoid function = 0.96, *p* < 10^−4^; I: R^2^ of fit for the sigmoid function = 0.98, *p* < 10^−4^). The black dashed line indicates the chance level. **H** Number of face-selective neurons. **I** Performance on the face classification task using only face-selective neurons.

First, we found that face-selective neurons are also observed in the randomized AlexNet with weight variation equivalent to pre-trained conditions (**Fig. 4B**, face). Furthermore, the observed neurons showed face-selective tuning such that the average selectivity index was significantly higher than that of the control as measured from the shuffled response (**Fig. 4C**, ***p < 0.001, Mann–Whitney U test). Similar to this face tuning, we also found that neurons spontaneously tuned to other non-face classes in this randomized AlexNet (**Fig. 4C**, non-face). However, the number of neurons responsive to non-face objects was significantly smaller than that of neurons responsive to face objects (**Fig. 4B**, ***p < 0.001, Mann– Whitney U test) and the average selectivity index of neurons responsive to non-face class was lower than that of face-selective neurons (**Fig. 4C**, ***p < 0.001, Mann–Whitney U test). These results imply that tuning for other non-face objects is not as strong as face-tuning at the population level.

Next, to test whether these spontaneous face-selective neurons enable classification of face images among other objects, we examined the SVM performance for face classification using the responses from the randomized AlexNet. We found that the SVM trained only with face-selective neurons shows performance comparable with that from using entire neurons in the final layer (face neurons only: 0.98 ± 0.02, All neurons: 0.99 ± 0.01). Furthermore, we confirmed that the SVM trained with face-selective neurons shows noticeably higher performance than that with neurons with no selectivity (**Fig. 4D**, *p < 0.05, Kolmogorov-Smirnov test).

Next, to investigate whether face-selectivity originated from a simple statistical variation of random feedforward projection, we reduced the weight variations of each kernel and examined changes in the face-selectivity of the neurons. We found that face-selective neurons respond less selectively to faces as the weight variation decreases (**Fig. 4E**). In addition, the face-like shapes of the preferred feature images in face-selective neurons were disrupted as the weight variation decreased (**Fig. 4F**). When the weight variation decreased to 53% of the original value, most neurons suddenly lost their face-selectivity (**Fig. 4G;** R^2^ of fit for the sigmoid function = 0.95; *p* < 10^−4^). Similarly, the number of face-selective neurons and the performance for the face classification task abruptly decreases when the weight variation is reduced to 53% of that in the pre-trained network (**Fig. 4H** and **4I;** R^2^ of fit for the sigmoid function = 0.96 and 0.98; *p* < 10^−4^ respectively). These results suggest that innate face-selective neurons can develop solely from the statistical variation present in the random initial wirings of bottom-up projections in the visual system, and that sufficient variation of the convolutional weights is critical to the spontaneous emergence of face-selectivity.

## Discussion

We showed that a biologically inspired DNN develops face-selective neurons without training, solely from statistical variation in the feedforward projections. These results suggest that the statistical complexity embedded in the structure of the neural circuit^33–35^ might be the origin of innate face-selective neuron development.

Our findings suggest a new scenario in which the proto-organization for face-selective neurons might be spontaneously generated, after which visual experience might sharpen and specify the selectivity of neurons. A recent fMRI study of the inferior temporal cortex in human infants and adult monkeys also supports this hypothesis^5^. Livingstone et al. show that the neurons broadly tuned to face are already observed in human infants (∼1 month old) and the region where these neurons were observed is identical to where the face-neurons of adult monkeys are observed. Furthermore, in the same study, it was reported that non-natural complex objects, such as man-made cars, provoke no selective neurons in infant monkeys^5^. This result implies that the innate template of face-selective neurons in infant monkeys may arise spontaneously and later may be fine-tuned during early visual experience. This is consistent with the results of this study.

State-of-the-art studies using random networks give important clues to how face-selective neurons could arise in our untrained model network. Recently, it was reported that an artificial network that learns visual features with random, untrained weights can perform image classification tasks^36,37^. Jarrett et al. showed that features from a randomly initialized one-layer convolutional network could classify the Caltech 101 dataset with performance level similar to that of a fine-tuned network, consistent with the mathematical notion that a combination of convolutional and pooling architecture could develop spatial frequency selectivity and translation invariance^38^. Overall, these results suggest that the initial structure of random networks might play important roles in visual feature extraction before the training process. It might even suggest that complex feature selectivity, such as face selectivity, might arise innately from the structure of the random feedforward circuitry.

It must be noted that the current results do not necessarily mean that spontaneous face-selectivity is the tuning observed in adult animals. There is plenty of evidence that the higher areas of the visual cortex are immature in the early development stage and that its functional circuit is modulated by visual experience^39–42^. There is also strong evidence that the IT region, where the face-selective neurons are observed, can be altered by early experience^43,44^. Considering anatomical and physiological changes that occur over the first postnatal year, the innate template of face-selective neurons in very early developmental stages must be refined by later visual experience including both bottom-up and top-down processes^45,46^. This scenario might be supported by recent observations of the existence of proto-organization of retinotopic organization and rough face-patch in higher regions of the visual cortex^5,47,48^. Moreover, observations on the early development of cortical circuits might provide further support to our scenario. Retino-thalamic feedforward projections are composed of noisy local samplings that result in un-refined receptive fields in individual thalamus neurons^49^. This is comparable to randomly initialized convolutional kernels before training. Spontaneous feature-selectivity generated in this early cortex might provide an initial basis for various visual functions and might be refined effectively when learning begins with visual experience.

In summary, we conclude that innate face-selectivity of neurons can spontaneously arise in a completely untrained neural network, solely from the statistical variance of feedforward projections. This finding suggests that various innate functions in the brain might originate from the organization of random initial wirings of the neural circuits, and provides new insight into the origin of innate cognitive functions.

## Methods

### Neural network model

We used the AlexNet^27^ as a representative model of the convolutional neural network. The network consists of feature extraction and classification networks. The feature extraction network consists of five convolutional layers with rectified linear unit (ReLU) activation and a pooling layer and the classification network has three fully-connected layers. The detailed parameters of the architecture drew upon *Krizhevsky et al*. (2012)^29^, which provided the models for V4 and IT^31^.

To figure out the origin of face-selective neurons, the three kinds of network were examined. (1) Pre-trained AlexNet: The network was trained to perform object recognition on the ILSVRC2012 ImageNet dataset. The network parameters, weights, and biases, were obtained using the MATLAB deep learning toolbox 2018b. (2) Permuted AlexNet: The network consists of randomly permuted weights and biases from those on each layer of the pre-trained AlexNet so that the spatial patterns in each kernel was disrupted by preserving the overall distribution. (3) Randomized AlexNet: The weights and biases of all the convolutional layers were initialized from Gaussian distributions with the same mean and the same (or reduced) standard deviation as those of each layer on the pre-trained AlexNet.

### Stimulus dataset

Two kinds of dataset were used. (1) Face vs non-face dataset: This set was used to find neurons that responded to face images selectively. A set of 100 face images and 1500 images of objects from 15 trained non-face classes of the pre-trained AlexNet (main classes: Animals, Vehicles, Fruits, Houses, Man-made objects). The non-face images were obtained from the ILSVRC2010 ImageNet dataset. Regarding the face class, the images were obtained from the VGGFace2 dataset, which consists of front-view faces of celebrities. (2) View-point dataset: This set was used to find neurons that invariantly responded to face images, even if imaged from different view-points. This dataset consists of 5 angle-based view-point classes (−90°, −45°, 0°, 45°, 90°) with 10 different Faces generated by the FaceGen Modeler Pro 3.18 (singular inversions) using the 10 center-view face images of different celebrities obtained from the VGGFace 2 dataset. For both datasets, the image size of the input to AlexNet was fixed at 227 × 227 pixels. Note that, according to the policy of bioRxiv that avoid the inclusion of photographs and any other identifying information of people, we show the illustration of faces instead of actual face pictures we used throuout this manuscript.

### Analysis of responses of the network neurons

The responses of the network neurons in the fifth convolutional layer were examined. Base on a previous study^26^, the face-selective neurons were defined as neurons that had significantly higher mean response to the face images than to the images of any non-face classes (p < 0.01, Mann–Whitney U test). To find non-face-class selective neurons, the same process was applied by replacing the face class with another one. To quantify the degree of tuning, a selectivity index of a single neuron to a preferred class was defined as the top-class selectivity referred from *Gale et al*. (2019)^32^. The selectivity index of a neuron was calculated as follows:

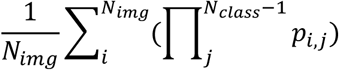

where *N*_*img*_ is the number of images on the preferred class (*N*_*img*_= 100), *N*_*class*_ is the number of all classes (*N*_*class*_ = 16) and *p*_*i,j*_ is the probability that the response of image *i* in the preferred class is greater than the response of all images in another class *j*. If the response of all images of the preferred class is larger than the response to images of all other classes, then the selectivity index is 1.

Among the Face-selective neurons found, a face view-point invariant neuron was defined as a neuron of which the response was not significantly different (p > 0.01, ANOVA) among all the view-point classes. The invariance index of a single neuron was set to one over the standard deviation of the view-point tuning curve, which was defined as the ratio of the mean to the variance of the mean response for each view-point class. Similar to the face-selective neurons, the view-point specific neurons were determined by the mean response of the preferred view-point class being significantly higher than that for any other view-point (p < 0.01, Mann–Whitney U test).

### Face vs non-face classification task for the network

A face vs non-face classification task was set to investigate whether face-selective neurons could perform basic face perception. To answer the question, an SVM was trained with network responses to images and predicted whether a class of unseen images was of face or not. To perform the task, the face vs non-face dataset was divided into a training and a test set (training set: test set = 2:1) with the same number of images in each class. Then, the label of the training set was changed to a binary class: face or non-face. For each trial, the SVM was trained with the relationship between the fifth layer’s response to the training set and to the new training label. After a training session, a model predicted a test label using the network response to the test set. To make a control case, the SVM was trained with a shuffled response to the training set to predict the test label.

### Receptive field analysis

To visualize the preferred input feature of single face-selective neurons, the receptive field was estimated by the reverse correlation method. The face and non-face image datasets (100 face images and 1,500 natural images of non-face objects) were used as stimuli. When the stimuli (N = 1,600) were presented on the network, the responses of the targeted neurons were measured. Every stimulus that generated response above the average response (all neurons to all images) was selected, weighted by corresponding response, and summed to obtain a preferred feature image. The receptive field was obtained by cropping the net input area of the target neuron on the preferred feature image. To measure the receptive field of other class selective or non-selective neurons, the same process was applied by replacing the targeted neuron.

---

**Supplementary Figure S1.**
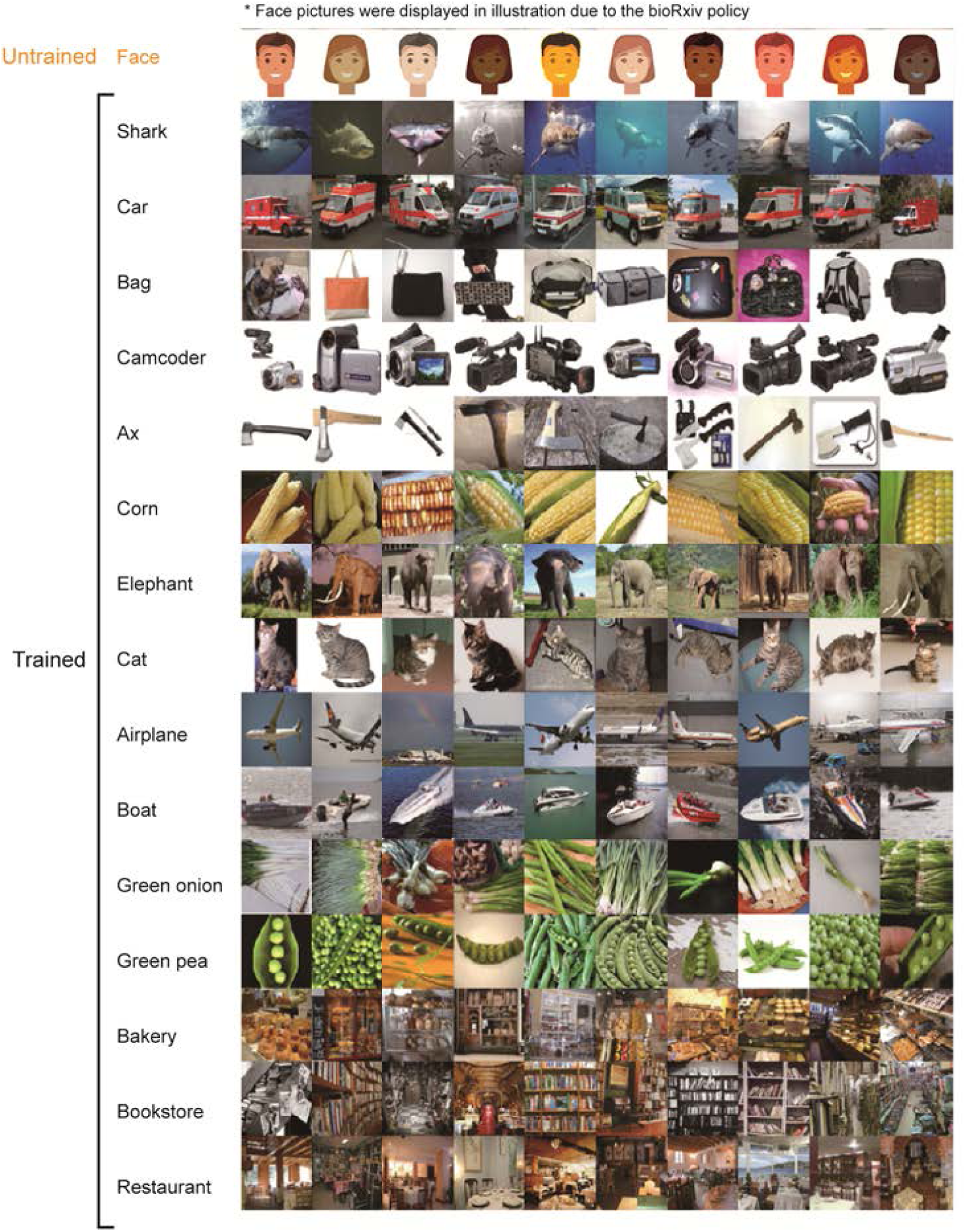
Image sets used for measuring selectivity in AlexNet. The image sets contain face and fifteen non-face image classes. Ten image samples were shown for each class. The non-face image samples were obtained from ImageNet dataset^50^.

**Supplementary Figure S2.**
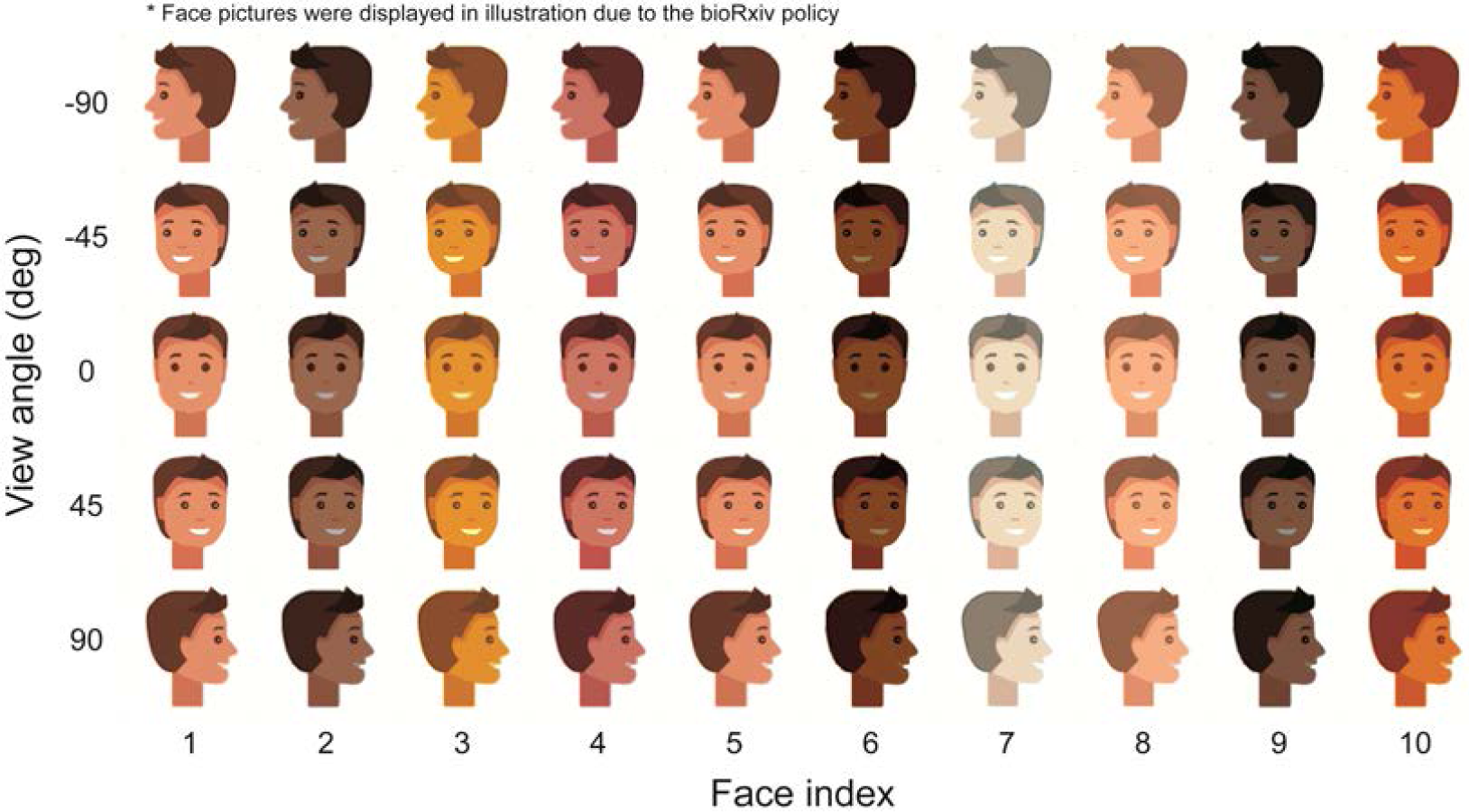
Image sets used for measuring invariance of face-selective neurons in AlexNet. Each of the face images, which was used to measure face-selectivity, was regenerated at five different angles from −90 to 90 degrees.

